# Biomass generation and heterologous isoprenoid milking from engineered microalgae grown in anaerobic membrane bioreactor effluent

**DOI:** 10.1101/2022.09.29.510234

**Authors:** Bárbara Bastos de Freitas, Sebastian Overmans, Julie Sanchez Medina, Pei-Ying Hong, Kyle J. Lauersen

## Abstract

1.

Wastewater (WW) treatment in anaerobic membrane bioreactors (AnMBR) is considered more sustainable than in their aerobic counterparts. However, outputs from AnMBR are mixed methane and carbon dioxide gas streams as well as ammonium- (N) and phosphate- (P) containing waters. Using AnMBR outputs as inputs for photoautotrophic algal cultivation can strip the CO_2_ and remove N and P from effluent which feed algal biomass generation. Recent advances in algal engineering have generated strains for concomitant high-value side product generation in addition to biomass, although only shown in heavily domesticated, lab-adapted strains. Here, investigated whether such a strain of *Chlamydomonas reinhardtii* could be grown directly in AnMBR effluent with CO_2_ at concentrations found in its off-gas. The domesticated strain was found to proliferate over bacteria in the non-sterile effluent, consume N and P to levels that meet general discharge or reuse limits, and tolerate cultivation in modelled (extreme) outdoor environmental conditions prevalent along the central Red Sea coast. High-value co-product milking was then demonstrated, up to 837 μg L^−1^ culture in 96 h, in addition to algal biomass production, ∼2.4 g CDW L^−1^ in 96 h, directly in effluents. This is the first demonstration of a combined bio-process that employs a heavily engineered algal strain to enhance the product generation potentials from AnMBR effluent treatment. This study shows it is possible to convert waste into value through use of engineered algae while also improve wastewater treatment economics through co-product generation.

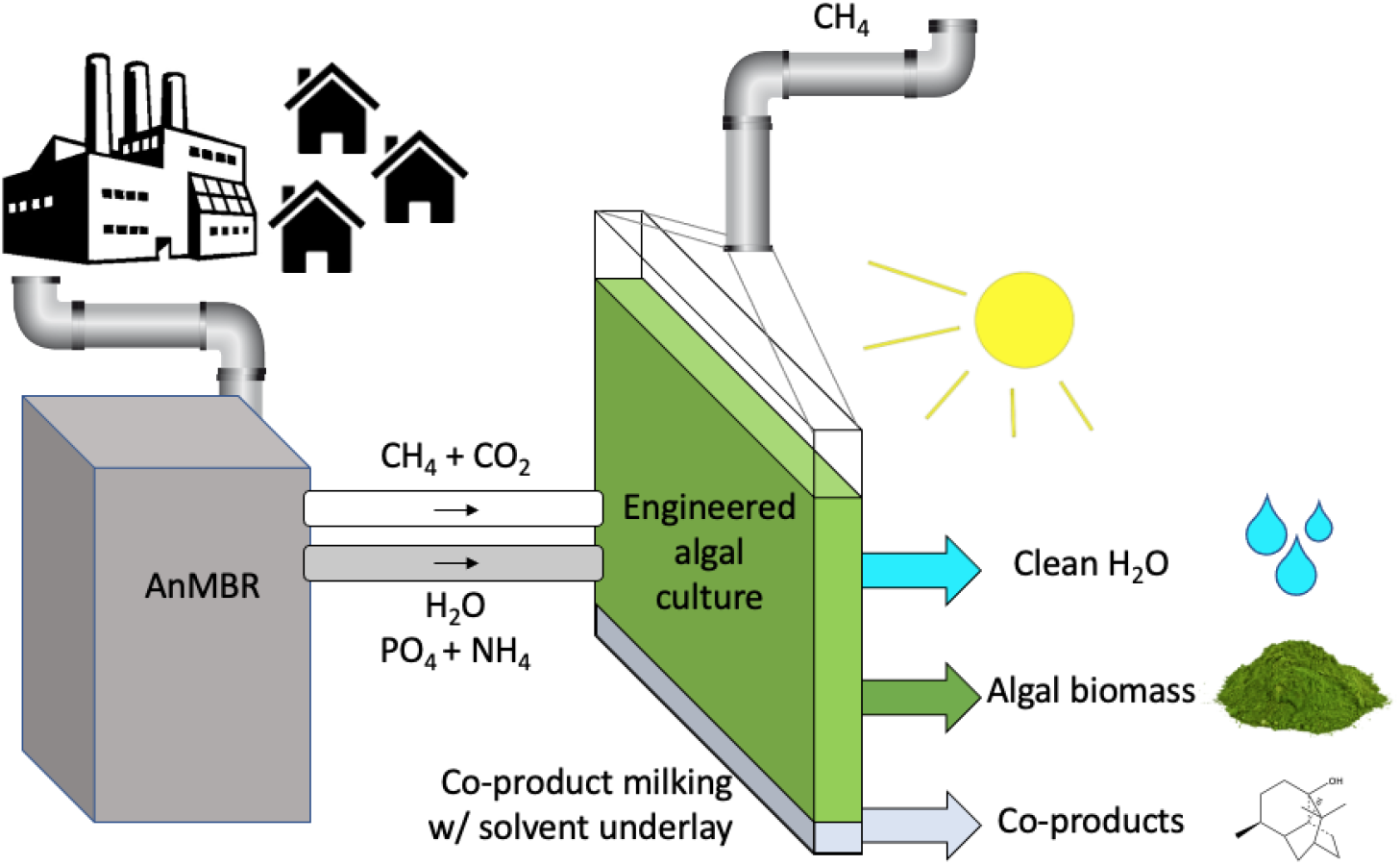

## 2. Introduction

Wastewater (WW) treatment strategies are an important part of human settlement infrastructure and an ongoing challenge of scale owing to the increasing human population (Cai et al., 2013; Liu and Hong, 2021). Conventional WW nutrient removal methods, both aerobic and chemical treatments, require high energy consumption and long process times while resulting in carbon emissions and excess sludge discharge (Li et al., 2019). In contrast, anaerobic membrane bioreactors (AnMBRs) have lower energy requirements while efficiently removing organic matter and suspended solids (Serna-García et al., 2020a). AnMBR process efficiencies are achieved by the combination of anaerobic microbial metabolism and high-surface area filtration technologies (Viruela et al., 2016). AnMBR technology yields biogas (methane, CH_4_) as a co-product of wastewater treatment, however, with a high percentage of carbon dioxide (CO_2_) which can be scrubbed to enrich methane for improved combustion. Ammonium and phosphate are also left in waters after AnMBR treatment, usually at levels that prohibit direct discharge into the environment (Gao et al., 2021; Xiong et al., 2018).

It has been proposed to couple AnMBRs with additional downstream cultivation of algae in photobioreactors to polish AnMBR effluents. Microalgae consume gaseous CO_2_ as a carbon source; ammonium and phosphate as nitrogen (N) and phosphorous (P) sources, respectively, by cellular processes driven by photosynthetic energy. During anaerobic digestion of wastewaters, existing organic N and P are converted into ammonium and inorganic phosphate, which algae are particularly adept at using as macronutrients (Serna-García et al., 2020b). Algal biomass can be a source of many valuable natural molecules, or itself be used as a feedstock for processes like the production of biochar or emerging bio-materials (Rajput et al., 2022). Due to their natural metabolic capacities, algae present promising biological systems for WW treatment process enhancement; excess nutrient and CO_2_ removal with concomitant (valuable) biomass generation (Cai et al., 2013; Chaudry, 2021; Leite et al., 2019; Viruela et al., 2016).

To increase the value and economics of algal bio-processes, engineering further levels of product generation into algal biomass through synthetic biology and metabolic engineering has been proposed. The model green microalga *Chlamydomonas reinhardtii* has now been extensively used to benchmark heterologous metabolite production from waste CO_2_ streams (Lauersen, 2019). Many examples of engineered co-product generation from *C. reinhardtii* have recently been shown: heterologous production of sesquiterpenes (Lauersen et al., 2016; Wichmann et al., 2018), diterpenes (Einhaus et al., 2022; Lauersen et al., 2018; Mehrshahi et al., 2020), polyamines (Freudenberg et al., 2022) and modified carotenoid pigments (Perozeni et al., 2020). Engineering efforts have been facilitated by the domestication of this alga through several rounds of mutation and transformation (Neupert et al., 2020, 2009) while advances in synthetic transgene design have enabled robust expression of heterologous pathways in this alga (Baier et al., 2020, 2018). The value of engineered algae is enhanced when the recombinant products can be harvested separately from the algal biomass using solvent-culture two-phase systems in a process called ‘microbial milking’ (Overmans et al., 2022; Overmans and Lauersen, 2022).

Highly domesticated and engineerable algae have only recently been reported, and bioprocesses that leverage these new organisms in practice have yet to be developed. Since these algae strains are adapted to laboratory conditions, they may not be as robust as algal species currently used in wastewater treatment or unsterile cultivation conditions. Here, we investigated the performance of highly domesticated *C. reinhardtii* strains directly in AnMBR effluent. We show that an engineered strain can simultaneously remove N and P from effluent, consume waste CO_2_, and yield the heterologous sesquiterpenoid patchoulol as co-product to biomass through metabolite milking. Our findings show that in addition to the valuable production of reclaimed water, co-products can be generated through engineered algae that can improve waste-stream treatment economics.

## 3. Materials and Methods

### 3.1. Effluent dilution experiments

*Chlamydomonas reinhardtii* strain UVM4 was graciously provided by Prof. Dr. Ralph Bock (Max Planck Institute of Molecular Physiology, Germany) under material transfer agreement to King Abdullah University of Science and Technology (KAUST). This strain is derived from several rounds of mutation of *C. reinhardtii* CC-4350 (Neupert et al., 2009). UVM4 contains a mutation in Sir2-type histone deacetylase (SRTA) that has been shown to improve transgene expression from the algal nuclear genome (Neupert et al., 2020). A recently reported derivative strain, which produces patchoulol was used in cultivations (Abdallah et al., 2022).

All algal cultures were maintained routinely on Tris acetate phosphate (TAP) or phosphite (TAPhi, Abdallah et al., 2022) agar plates under 50 μmol photons m^−2^ s^−1^ light intensity before being transferred into 45 mL medium in 125 mL Erlenmeyer flasks with liquid medium and shaken at 120 rpm on a 12 h:12 h dark:light (150 μmol photons m^−2^ s^−1^) cycle. Light spectra of different illumination set ups for all cultivations are reported in Suppl. Data 1. After preculturing in shake flasks, 45 mL of cells were centrifuged at 1000 x g for 5 min and resuspended in 400 mL AnMBR effluent, allowed to establish as culture for 3 days and then further diluted into effluent in 400 mL photobioreactor flasks.

Cultivations were performed in Algem photobioreactors (Algenuity, UK) at four different dilutions of raw AnMBR effluent (100%, 75%, 50%, and 25%) in double distilled water in biological duplicates. The strain was grown at 25 ºC with a 7% CO_2_ air mixture in 400 mL volumes, with a starting algal cell concentration of 3×10^5^ cells mL^−1^, shaken at 100 rpm, and a 12 h:12 h dark:light cycle, with a gradual increase from 0 to 325 μmol photons m^−2^ s^−1^ for 144 h (Fig. 1a). A gas mix of 7% CO_2_ in ambient air was supplied periodically to cultures at 25 cc min^−1^ according to the pH control level set on the reactor. Optical densities were measured by the photobioreactors in 10 min intervals and subsequently converted to algae biomass (g L^−1^) using a standard curve made using dried biomass samples (Suppl. Data 2). The raw AnMBR effluent contained approximately 20 mg L^-1^ ammonium (NH_4_^+^-N) and 10 mg L^-1^ phosphate (PO_4_^3-^-P), as measured by HACH kits (see section 2.5).

**Figure 1.**
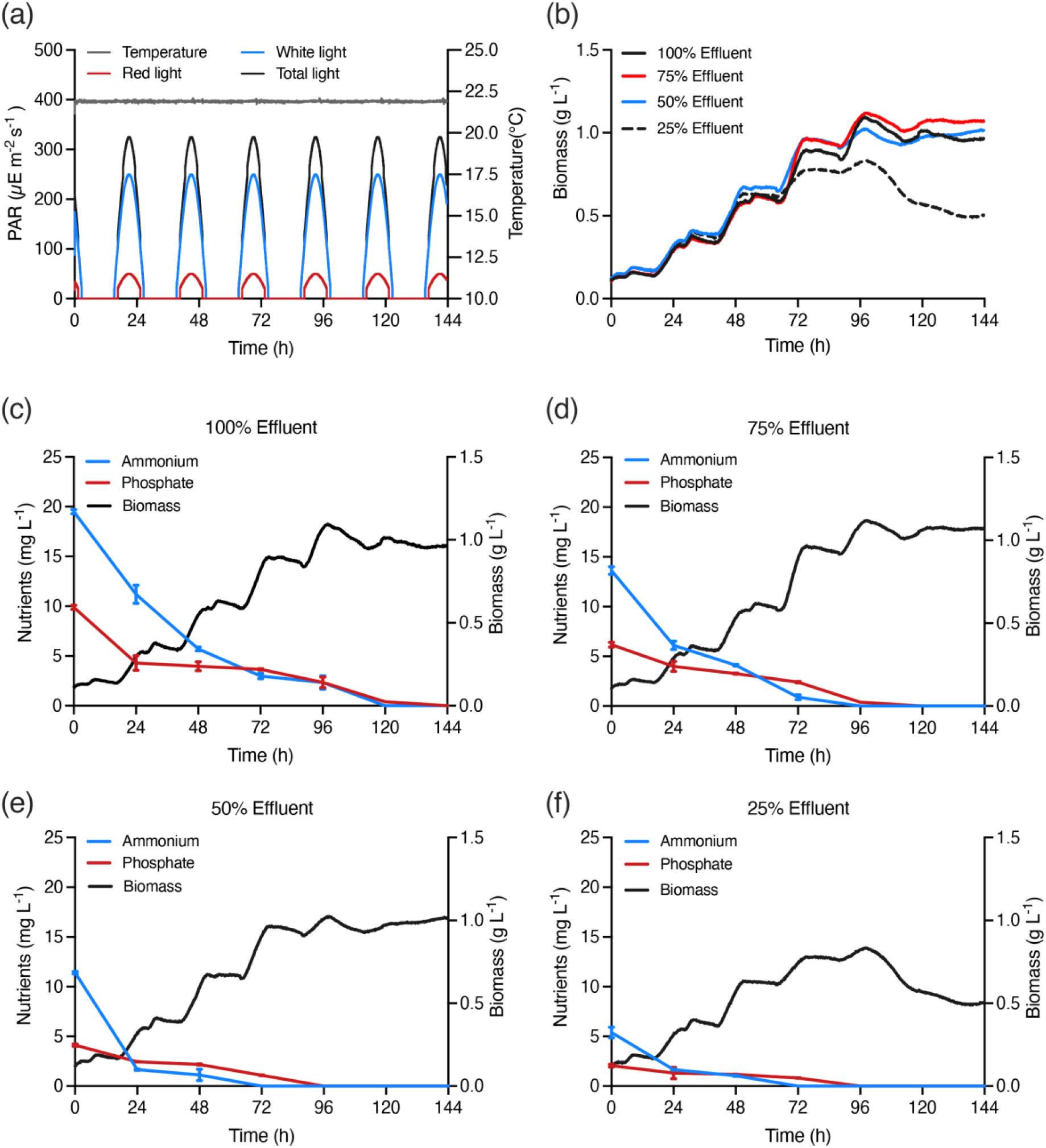
(a) Temperature and light conditions and (b) biomass generated at different effluent percentages during the effluent dilution experiment. Lower panels show ammonium and phosphate concentrations over time when 100% effluent (c), 75% effluent (d), 50% effluent (e) and 25% effluent (f) dilution were used as growth medium.

### 3.2. Local weather condition simulations and algal culture performance

We tested the feasibility of cultivating the domesticated UVM4 strain in raw AnMBR effluent using outdoor light and temperature conditions of Thuwal, Saudi Arabia (22.3046N, 39.1022E) by weather simulation in photobioreactors. The weather data used for this experiment consisted of temperature and photosynthetically active radiation (PAR) measurements (10 min intervals) recorded at the Coastal and Marine Resources Core Lab (CMR) at KAUST between January and December 2014 (Suppl. Data 3). Twelve 400 mL *C. reinhardtii* UVM4 cultures in raw AnMBR effluent with a starting cell concentration of 3×10^5^ cells mL^−1^ were grown in Algem photobioreactors under different temperature and light conditions, simulating the months January– December 2014. Cultures were shaken at 100 rpm for 96 h, with 7% CO_2_ injections supplied periodically for pH control, as described above. The optical densities were measured in 10 min intervals and subsequently converted to algae biomass (g L^−1^) as above (see Suppl. Data 4).

### 3.3. Waste stream capture and concomitant patchoulol production

A previously engineered *C. reinhardtii* transformant line UPN22-2x*Pc*Ps, which produces the heterologous terpenoid alcohol patchoulol, was used in this experiment (Lauersen et al., 2016; Neupert et al., 2009). The algal culture was maintained on TAPhi NO_3_ agar plates, transferred into small volumes of the same liquid medium, and subsequently grown in Erlenmeyer flasks under ∼100 μmol photons m^-2^ s^-1^ and 120 rpm shaking before bioreactor cultivation. The pre-culture was then harvested by centrifugation at 1000 x g for 5 min, and the cells resuspended directly in AnMBR effluent with a starting cell concentration of 5×10^5^ cells mL^−1^.

This experiment was performed in biological triplicate, whereby 3×6 2L Erlenmeyer flasks were used to grow 400 mL cultures of the UPN22-2x*Pc*Ps strain in Algem photobioreactors. Per experiment, six cultures were grown in reactors shaken at 100 rpm for 96 h, with CO_2_ injections supplied periodically for pH control, as described above. Each replicate was grown under one of six experimental conditions as described below: constant light, 12 h:12 h light:dark, both at 25ºC and 325 μmol photons m^-2^ s^-1^ as controls, and 4 different temperature and light conditions that each represent one of the four seasons in Thuwal, Saudi Arabia. Local weather station data were used to model 1 week from each of February (Winter), May (Spring), August (Summer), and November (Autumn) from 2014 (Suppl. Data 2) in the reactors. To capture patchoulol produced from the algal cells during growth, 20 mL of the perfluorinated amine FC-3283 (Acros Organics, Geel, Belgium) was added to each culture as an underlay, as recently described (Overmans and Lauersen, 2022).

Daily samples of 10 mL algae culture from each replicate flask were taken for determination of bacterial- / algal-cell concentrations, biomass quantification, and nutrient analysis (Suppl. Data 5), while 500 μL of FC-3283 was sampled daily from each replicate bioreactor flask to quantify the accumulation of patchoulol product in the underlay. FC-3283 samples were stored at -20°C until the end of the experiment when all samples were together processed for GC-MS analysis.

### 3.4. Culture growth measurements

Algal culture growth was determined by measuring cell densities with an Invitrogen Attune NxT flow cytometer (Thermo Fisher Scientific, UK) equipped with a Cytkick microtiter plate autosampler unit. Prior to analysis, each biological replicate sample was diluted 1:100 with 0.9% NaCl solution. Of each diluted sample, 250 μL was measured in technical duplicates (n=2) using a 96-well microtiter plate loaded into the autosampler. Samples were mixed three times immediately before analysis, and the first 25 μL of sample was discarded to ensure a stable cell flow rate during measurement. Data acquisition was stopped when 50 μL from each well was analyzed. All post-acquisition analyses and population clustering were performed using Attune NxT Software v3.2.1 (Life Technologies, USA).

As raw effluent was used as a culture medium in unsterile conditions, we also determined bacterial cell counts using the same flow cytometer: 80 μL of each culture sample was diluted in 720 μL of 1× PBS, then stained with 8.08 μL of 1X Invitrogen SYBR Green nucleic acid stain (Thermo Fisher Scientific, Carlsbad, CA, USA) and incubated for 10 min at 37 °C before measurement.

Algae biomass was measured by comparing it to a standard curve established using *C. reinhardtii* UVM4 and 2xPcPs cultures to determine the corresponding dry weight. These curves were obtained by measuring absorbance at 754 nm using a spectrophotometer (Genesys 10S UV-VIS, Thermo Fisher Scientific, UK) and correlating the relative optical density to dry biomass.

### 3.5. Nutrient analysis

To analyse N and P concentrations in culture medium at different time points, 10 mL of each algae culture in wastewater was sampled daily, centrifuged at 4500 x g for 5 min at 20 °C, and the supernatants collected. Prior to analysis, all supernatants were filtered with a 0.45 μm syringe filter to remove insoluble particles. Ammonium (NH_4_^+^-N) and phosphate (PO_4_^3-^-P) concentrations were determined spectrophotometrically with a DR 1900 spectrophotometer (Hach, Germany) after the samples were prepared with AmVer High Range and TNT 844 analysis kits (Hach, Loveland, Colorado, USA), respectively, following the manufacturer’s protocols. Each nutrient concentration was analysed in technical duplicates (n = 2).

### 3.6. Gas chromatography

The daily collected FC-3283 samples were analyzed using an Agilent 7890A gas chromatograph (GC) that was equipped with a DB-5MS column (Agilent J&W, USA). The instrument was attached to a 5975C mass spectrometer (MS) with a triple-axis detector (Agilent Technologies, USA). We used a previously described GC oven temperature protocol (Overmans and Lauersen, 2022). All GC-MS measurements were performed in technical duplicates (n=2). Chromatograms were manually reviewed for quality control before patchoulol GC peak areas were integrated using MassHunter Workstation v. B.08.00 (Agilent Technologies, USA). A patchoulol standard (18450, Cayman Chemical Company, USA) five-point calibration curve in the range of 6.25–50 μg patchoulol mL^−1^ in FC-3283 was used for product quantification (see Suppl. Data 6).

### 3.7. CO_2_ capture calculations

Theoretical CO_2_ capture (g L^−1^) was calculated using measured algal biomass (g) values as shown in Eqn 1:

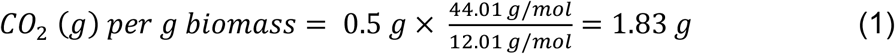

Where 0.5 g represents the previously reported approximate mass of carbon per gram of biomass (Chisti, 2007), while 44.01 g/mol and 12.01 g/mol refer to the molecular masses of CO_2_ and carbon, respectively.

## 4. Results and Discussion

### 4.1. *C. reinhardtii* UVM4 cultivation in raw AnMBR effluent

*C. reinhardtii* UVM4 was cultivated in different dilutions of raw AnMBR effluent to evaluate the growth of this strain in non-sterile conditions with the inherent N and P of this liquid. AnMBR effluent contains ammonium and phosphate, the C:N and C:P ratios are generally low and support autotrophic algal growth more than heterotrophic bacterial growth. Cultures were sparged with clean 7% CO_2_ to provide a carbon source similar to what would come from an AnMBR reactor and to regulate culture pH. As algae grow, they consume CO_2_, which raises culture pH. CO_2_ injection for pH control in this way minimizes wasted CO_2_ as it is only supplied as the algae consume it.

Other green algae like *Scenedesmus* sp. and *Chlorella* sp. have both been shown to grow well in AnMBR effluent (Serna-García et al., 2020b). A recent *Chlamydomonas* sp. isolate from WW has been shown to grow robustly in non-sterile WW conditions and can outcompete bacterial contaminants (Klassen et al., 2020). However, few studies have shown *C. reinhardtii* growth for WW post-treatment, fewer for effluent from AnMBRs, and none to date have used a heavily domesticated and lab-adapted genetically engineered strain. *C. reinhardtii* UVM4 has been derived from several rounds of mutagenesis and transformation of the parent wild-type strain (Neupert et al., 2009). Given that this strain lacks a cell wall, has been long-adapted to nutrient surplus conditions, and may be outcompeted by other organisms, it was important to determine its ability to grow directly in unsterile AnMBR effluent.

Mixed microalgae-bacteria photo-bioreactors are able to remove nutrients and CO_2_ from AnMBR effluent (Xiong et al., 2018), and the growth of *C. reinhardtii* UVM4 on either pure or diluted effluent was demonstrated here (Fig. 1b). The strain grew directly in non-sterile effluent and completely consumed N and P as nutrients for its biomass after 144 h cultivation (Fig. 1c). *C. reinhardtii* UVM4 was found to grow in all dilutions of effluent in water, with only the 25% dilution exhibiting reduced growth behaviours (Fig. 1b). The effluents used for this experiment initially contained 23–25 mg L^-1^ of ammonium (NH_4_^+^-N) and 7–8 mg L^-1^ of phosphate (PO_4_^3-^-P). The raw effluent (100%) and 50% dilution were suitable substrates for algal culture growth, with nutrient removal occurring earlier in the 50% dilutions (Fig. 1c and e). Previous microalgae-bacteria cultures have been shown to have excellent stability as the symbiosis of microalgae and bacteria can even improve pollutant removal in wastewater (Liu and Hong, 2021). Mixed cultures have also been shown to promote growth of other *Chlamydomonas* spp. (Klassen et al., 2020). Here, the heavily domesticated *C. reinhardtii* UVM4 was robust enough to be the dominant microorganism in the AnMBR effluent under non-sterile conditions (Suppl. Fig. 1) and yield water which had nutrient levels acceptable for discharge (Al-Jasser, 2011). This finding encouraged further investigation into how an engineered alga could be used for post-treatment of AnMBR effluent.

### 4.2. *C. reinhardtii* growth under extreme climate with raw AnMBR effluent

The ability of *C. reinhardtii* UVM4 to grow in raw AnMBR effluent in simulated weather conditions (temperature and PAR) from the Saudi Arabian Red Sea shores was tested (Fig. 2 a). The Saudi Arabian central Red Sea is characterized by an arid climate, with daytime summer air temperatures of > 40°C(Almazroui et al., 2012; Alsarmi and Washington, 2014). In addition, the urban population of Saudi Arabia is growing, and effective wastewater treatment strategies are required that are compatible with the local environmental conditions. Here, domesticated *C. reinhardtii* cultures grown directly in AnMBR effluent were found to tolerate the local light and temperature regime across the year, with slightly lower biomass generated under the winter conditions (Fig. 2 b).

**Figure 2.**
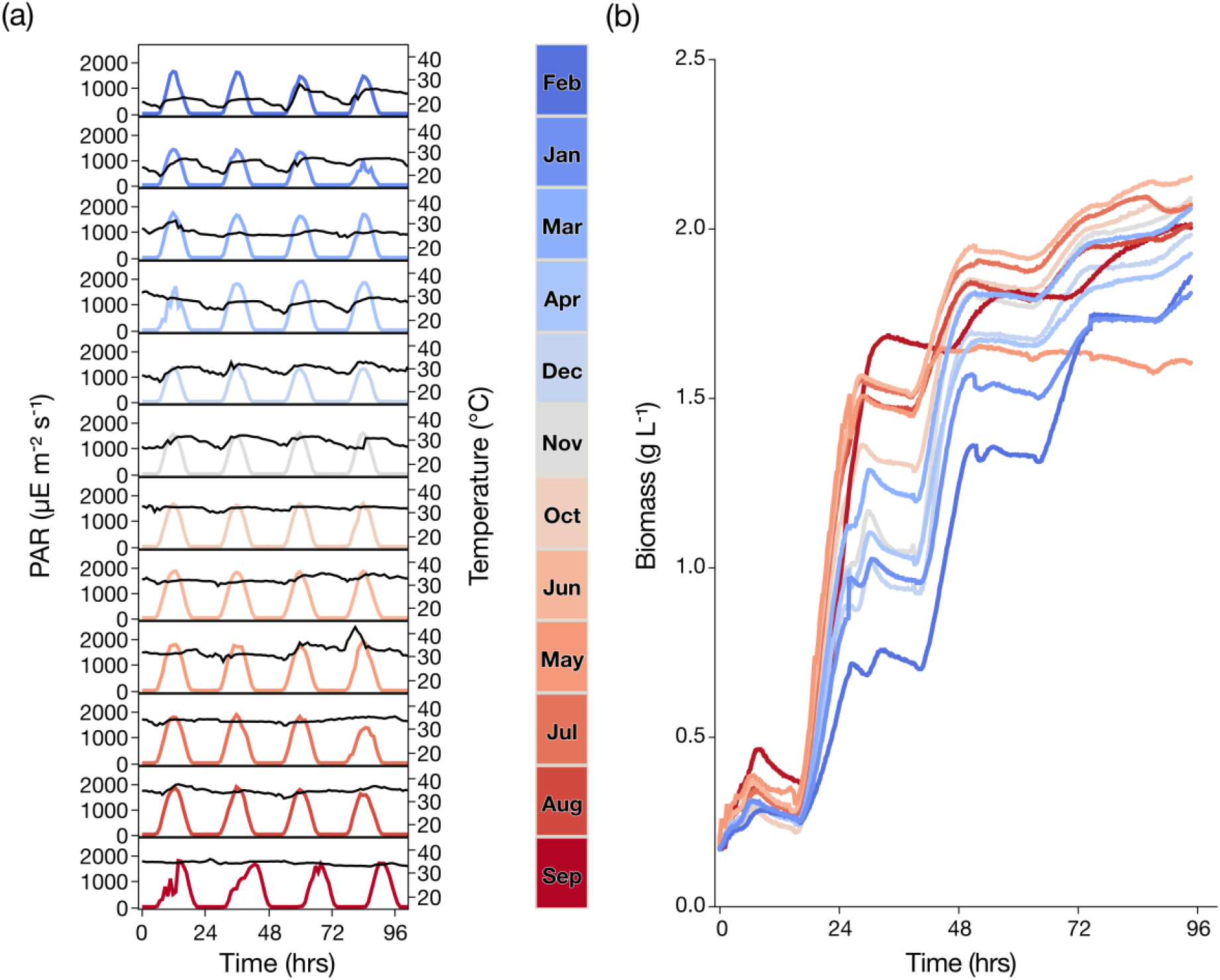
(a) Temperature and PAR profiles that were recorded in Thuwal, Saudi Arabia between January and December 2014. Displayed are the first 96h of each month, which were used for the local weather simulation experiment. The black lines represent temperature, while the colored lines represent PAR. (b) Algal biomass accumulated over 96 h under the different simulated local temperature-PAR conditions shown in (a).

During most modelled months, the algal biomass of AnMBR effluent batch cultivation was ∼2g L^−1^ by 96 hours (Fig. 2 b). The highest biomass production was observed in the modelled month of June (2.26 g L^-1^, 29–35°C, Fig. 2 a). During August, even higher temperatures are observed at this locale (T: 31–37°C). Nevertheless, the UVM4 strain grew well, reaching 2.12 g CDW L^-1^ (Fig. 2 b). Bacterial contamination was present in all conditions but did not cause algal culture crash, nor did bacteria outcompete the culture during the investigation period (Suppl. Fig. 2).

Environmental modelling using local weather data in replicate bioreactors, as shown here, will be a valuable tool for benchmarking strain performance for outdoor cultivation in different locales. This is the first example of such modelling for an engineered *C. reinhardtii* strain grown in WW, and complements previous efforts to grow engineered algae and cyanobacteria in outdoor conditions (Wichmann et al., 2021). Full-year modelling with each month provided valuable insight into how processes may perform in the local context throughout the year. Conditions from specific months were then chosen to represent seasonal changes to model outdoor bio-process performance using a strain engineered to produce the heterologous sesquiterpenoid patchoulol, and assess its product yields when grown directly in AnMBR effluent.

### 4.3. Specialty chemical co-production during effluent nutrient removal by an engineered green alga

UVM4 has previously been genetically and metabolically modified to produce the heterologous plant sesquiterpenoid patchouli alcohol, “patchoulol” (Abdallah et al., 2022; Lauersen et al., 2016). Heterologous terpenoid products can be collected during cultivation of microbial cultures using a two-phase culture-solvent system (Beekwilder et al., 2014; Lauersen, 2019; Overmans et al., 2022). The hydrophobic terpenoid products accumulate in biocompatible solvents as the products are more favored to partition into the solvents than into the cells or culture medium (Gruchattka and Kayser, 2015). Recently, we have shown a novel approach to this continuous culture-solvent interaction, or ‘milking’, by employing perfluorinated liquids that form a layer underneath the aqueous culture medium, an ‘underlay’, due to their higher densities (Overmans and Lauersen, 2022). These liquid fluorocarbons (FCs) are inert and accumulate terpenoid products from microbial cultures efficiently. Unlike organic solvent overlays, FCs can be effectively used in the Algem photobioreactors, due to their high density and physical dynamics, to capture terpenoid products during algal cultivation.

The engineered, patchoulol-producing *C. reinhardtii* was cultivated in representative seasonal conditions (Spring, Summer, Autumn, Winter) as well as in two artificial illumination conditions: continuous light and a 12 h:12 h light:dark cycle at constant temperature (Fig. 3). Artificial conditions were used to compare controlled culture conditions with ‘outdoor’ conditions that could be employed for algal-based AnMBR effluent nutrient removal at large scales as in the case of municipal WW treatment. The patchoulol-producing *C. reinhardtii* was found to grow well in AnMBR effluent, generating 2.35, 2.18, and 2.43 g CDW L^−1^ culture in May, August, and November conditions, respectively, after 36–60 h of cultivation (Fig. 3b). Continuous light generated 2.7 g CDW L^−1^ after 60 h of cultivation, whereas in February and the 12 h:12 h light conditions, biomass reached a maximum of ∼2.0 g CDW L^−1^ after 80 h (Fig. 3b).

**Figure 3.**
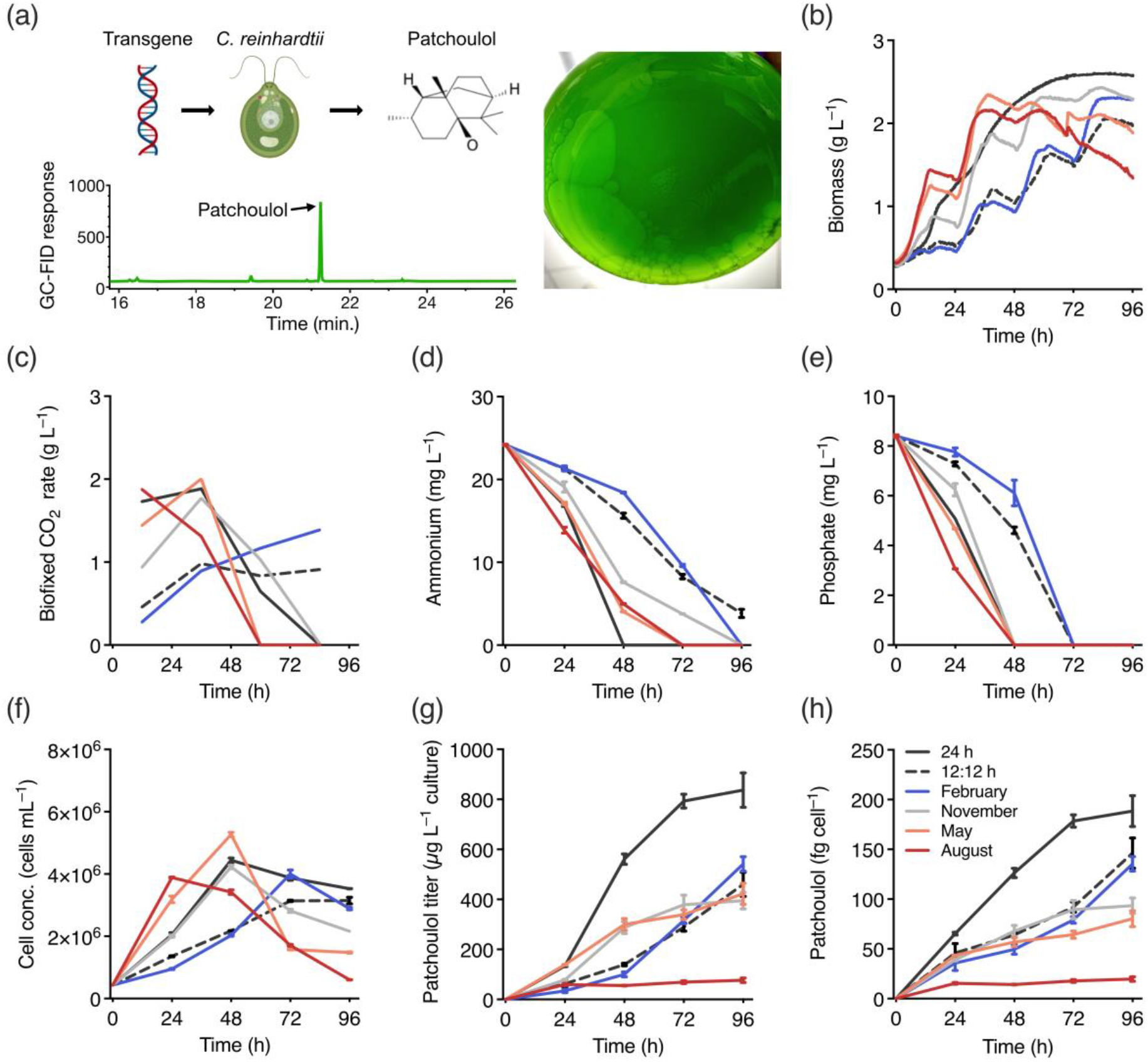
(a) Schematic of transgene insertion into *C. reinhardtii* which leads to patchoulol production, a photo showing the FC-3283 underlay in a bioreactor flask, and a representative GC-MS chromatogram of an FC-3283 sample containing patchoulol (b) algal biomass (c) bio-fixed CO_2_, calculated as the maximum biomass in a 24 h interval * 1.83 (CO_2_ : biomass ratio) (d, e) ammonium and phosphate concentrations and (f) algal cell counts during 96 h of *C. reinhardtii* cultivation in wastewater in six different environmental conditions. (g, h) Volumetric and cell-dependent patchoulol production during cultivation. Values displayed in (d)–(h) are means ± SEM of two technical replicates (n=2).

Across all conditions, patchoulol production gradually increased over the 96 h experimental period (Fig. 3g–h; Suppl. Fig 3). Continuous light yielded the highest patchoulol production, with a maximum titer of 837 μg patchoulol L^−1^ culture after 96 h (Fig. 3g). Under the 12 h:12 h light:dark illumination cycle, and the simulated February, November and May conditions, patchoulol titers were relatively similar to each other by the end of the experiment, with concentrations of 460, 542, 395 and 423 μg L^−1^ culture. Under August conditions, patchoulol production was considerably lower (76 μg L^−1^ culture). Patchoulol production per cell, based on algal cell count measurements (Fig. 3 f), showed a similar pattern to the volumetric yields. Cell-specific patchoulol production was highest in the continuous light regime (188 fg cell^−1^) and negligible in the summer conditions (August: 20 fg cell^−1^) (Fig. 3h). Moderate production of patchoulol per-cell was observed under the 12 h:12 h light:dark illumination (146 fg cell^−1^), the simulated February (136 fg cell^−1^), November (93 fg cell^−1^) and May (80 fg cell^−1^) conditions. The latter finding suggests that patchoulol production per cell gradually decreases with increasing cultivation temperature.

### 4.4. Culture of engineered alga in unsterile effluent and biomass accumulation

In this experiment, highly engineered algal biomass was also dominant in these unsterile cultivations, despite the presence of contaminating bacteria (Suppl. Fig. 4. Algal-bacterial consortia in wastewater treatment simultaneously degrade organic carbon by heterotrophic bacteria and nutrients (N and P) by photoautotrophic algae. In their interactions, the oxygen produced by algae sustains bacterial respiration that contributes to nitrification and denitrification as well as organic matter removal. In return, the CO_2_ produced by bacteria enhances microalgae growth (Foladori et al., 2018; Gou et al., 2020). Qu et al. (2020) reported that *Chlamydomonas* sp. QWY37 remediated swine wastewater and reduced the amount of total N by 96% and removed 100% of P. A newly isolated strain, named *Chlamydomonas* sp. YC, exhibits high tolerance towards ammonium, and was used to treat the undiluted rare earth element mining wastewater (Zhou et al., 2022). Here, the engineered *C. reinhardtii* patchoulol-producing strain removed all ammonium and phosphate from the raw effluent by the end of the cultivations (96 h), except for the NH_4_^+^-N content in the 12 h:12 h dark:light cultures (Fig. 3 d–e, and Suppl. Fig. 5). It could be expected that some cell wall proteins of the alga may be present at the end of cultivation, which would be removed by any ultrafiltration steps used for biomass concentration. Under continuous light and conditions simulating May and August, the same strain was able to take up all the NH_4_^+^-N after only 72 h and phosphates after 48 h (Fig 3 d–e). As pH was controlled (7–7.1) across experimental conditions, NH_4_^+^-N volatilization and PO_4_^3-^-P precipitation were not issues here, unlike in previous studies (Gutzeit et al., 2005).

It was observed that algal biomass generation was highest in May and August conditions after 24–48 h (Fig. 3 b), whereas the rate of biomass generation decreased after the consumption of N and P sources (Fig 3 d–e). These findings point to potential repetitive batch scheduling conditions, which could be used to cycle nutrient removal with biomass generation.

One of the advantages of using microalgae to treat WW is their photosynthetic CO_2_ fixation, which can also be used to assist the cleaning of CO_2_ from CH_4_ in AnMBR off-gas (Ruiz-Martinez et al., 2012). A previous report also found that CO_2_ from a lab-scale AnMBR was completely consumed by algal culture, but only after 4 d of operation (Xiong et al., 2018). Here, 7% CO_2_ without CH_4_ was provided as a carbon source and pH control in order to see maximal growth of algae with the available N and P of the effluent. Our cultivations yielded CDW of over 2 g L^−1^ by 48h in the simulated August, May, November, and continuous culture conditions (Fig. 3b). For May and continuous illumination conditions, rates of CO_2_ to biomass (biofixation) were ∼ 2 and 1.88 g L^−1^, respectively, in the 24 hours of the second day of cultivation (Fig. 3c). The growth performance of the strain and biomass accumulation over 96 hours are equivalent to ∼3.66 and 3.44g of CO_2_ into biomass L^−1^ algal culture, for May and August, respectively. The AnMBR used here generates 631.5 mg of CO_2_ L^−1^ effluent. It should then be possible for the engineered *C. reinhardtii* strain to capture 100% of the CO_2_ generated in its off-gas using the N and P found in 1 L effluent in 24 hours of the logarithmic growth phase of these conditions. However, cultivation periods longer than 24 hours would be required for complete N and P consumption as this took 48-72 hours in the conditions tested here (Fig. 3 d,e). Therefore, AnMBR off-gas storage and controlled feeding to algal cultures would need to be independently controlled to maximize N and P removal from the effluent, while scrubbing CO_2_ from the gas mixture. These process factors would need to be considered in bio-process design scheduling.

The high temperature and light conditions typical for the central Red Sea coast of Saudi Arabia, seem to be favorable for rapid growth of the engineered *C. reinhardtii* strain even in this effluent. The milking of a terpenoid co-product during nutrient removal could be considered a highly promising way to increase the economics of lost resources in WWs. Solvent milking can be cyclic, wherein the extraction of the patchoulol from the solvent can be performed and the solvent returned to culture in low OPEX bio-processes (Overmans et al., 2022). Here, we chose the sesquiterpenoid patchoulol as an exemplary product of the engineered algae due to the availability of strains. Different terpenoids produced by engineered algal could be chosen based on market demands as many of these products have been shown and engineering is ongoing (Lauersen, 2019). The market price of synthetic patchoulol made by engineered yeast is ∼$1.2 USD g^-1^ (Clearwood™ by Firmenich https://perfumersupplyhouse.com/product/clearwood-firmenich/). Patchoulol productivity from the engineered algae was up to 837 μg L^-1^ algal culture in effluent (96 h) and between 200-400 μg L^-1^ culture d^-1^ across biological replicates in the highest growth phase of 24-48 h (Fig. 3g, Supp. Fig. 3). At these productivities, $1.2 USD could be generated for every 1,975 L (96 h batch) or 5,000-2,500 L d^-1^ (high growth-rate repetitive batch) of effluent.

Higher temperatures in August simulations resulted in drastically lower patchoulol co-product yields, indicating unfavorable isoprenoid metabolism in these higher temperatures despite high biomass production rates (Fig. 3). In other conditions with lower temperatures, patchoulol production was higher although nutrient removal and algal growth were slower. Therefore, a balance in process design will perhaps be necessary wherein seasonal outdoor, lower energy input, processes are switched with semi artificial cultivation conditions in summer conditions. These factors for co-product generation will also need to be balanced with culture residence times to ensure complete N and P removal, gas-stream CO_2_ stripping and biomass separation.

## 5. Conclusions

We show here that a highly domesticated, lab-adapted and engineered green alga can grow directly in unsterile AnMBR effluents and yield considerable biomass (2.7 g CDW L^−1^); even under extreme light and temperature conditions typical for the central Red Sea coast of the Arabian Peninsula. Using a co-product milking strategy, we could simultaneously generate clean water and algal biomass, and convert waste CO_2_ into a valuable co-product patchoulol, which was non-intrusively collected through microbial milking. This process could readily be extended to other high-value isoprenoids, secreted recombinant proteins, or volatile bulk chemical feedstocks produced by engineered algae. The biomass itself is valuable, although engineering alternative pigments could further enhance this. Alternatively, algal biomass generated through post-treatment of AnMBR effluent could be used as a bulk vehicle for carbon sequestration, and nutrient recovery strategies, such as biomass conversion to biochar, hydrothermal liquefaction, conversion to asphalt, generation of phytostimulant fertilizers, or direct use in bio-materials (plastics) production. The demonstration that domesticated and engineerable algal strains can grow directly in AnMBR effluent and yield high value side products drastically expands the range of possibilities to increase the circular value of these systems. This can potentially be done by having municipal wastewater go through AnMBR-based treatment, and the AnMBR effluent being fed into a photobioreactor inoculated with the engineered algal strains to yield valuable products and/or biomass.

## Supporting information

Supplemental files zip

## 6. Author Contributions

BF and SO were responsible for design and performed experiments and contributed to writing the manuscript. JM operated the AnMBR and sourced the effluent. PYH and KJL contributed to experimental design, project supervision, funding acquisition, and manuscript writing.

## 8. Acknowledgements

We would like to express special thanks to the Coastal and Marine Resources Core Lab (CMR) at KAUST for providing the temperature and PAR data sets used in this study.

## References

Abdallah, M.N., Wellman, G.B., Overmans, S., Lauersen, K.J., 2022. Combinatorial Engineering Enables Photoautotrophic Growth in High Cell Density Phosphite-Buffered Media to Support Engineered Chlamydomonas reinhardtii Bio-Production Concepts. Front Microbiol 13, 1–12. https://doi.org/10.3389/fmicb.2022.885840

Al-Jasser, A.O., 2011. Saudi wastewater reuse standards for agricultural irrigation: Riyadh treatment plants effluent compliance. Journal of King Saud University - Engineering Sciences 23, 1–8. https://doi.org/10.1016/j.jksues.2009.06.001

Almazroui, M., Nazrul Islam, M., Athar, H., Jones, P.D., Rahman, M.A., 2012. Recent climate change in the Arabian Peninsula: Annual rainfall and temperature analysis of Saudi Arabia for 1978-2009. International Journal of Climatology 32, 953–966. https://doi.org/10.1002/joc.3446

Alsarmi, S.H., Washington, R., 2014. Changes in climate extremes in the Arabian Peninsula: Analysis of daily data. International Journal of Climatology 34, 1329–1345. https://doi.org/10.1002/joc.3772

Baier, T., Jacobebbinghaus, N., Einhaus, A., Lauersen, K.J., Kruse, O., 2020. Introns mediate post-Transcriptional enhancement of nuclear gene expression in the green microalga Chlamydomonas reinhardtii. PLoS Genet 16, 1–21. https://doi.org/10.1371/journal.pgen.1008944

Baier, T., Wichmann, J., Kruse, O., Lauersen, K.J., 2018. Intron-containing algal transgenes mediate efficient recombinant gene expression in the green microalga Chlamydomonas reinhardtii. Nucleic Acids Res 46, 6909–6919. https://doi.org/10.1093/nar/gky532

Beekwilder, J., van Houwelingen, A., Cankar, K., van Dijk, A.D.J., de Jong, R.M., Stoopen, G., Bouwmeester, H., Achkar, J., Sonke, T., Bosch, D., 2014. Valencene synthase from the heartwood of Nootka cypress (Callitropsis nootkatensis) for biotechnological production of valencene. Plant Biotechnol J 12, 174–182. https://doi.org/10.1111/pbi.12124

Cai, T., Park, S.Y., Li, Y., 2013. Nutrient recovery from wastewater streams by microalgae: Status and prospects. Renewable and Sustainable Energy Reviews 19, 360–369. https://doi.org/10.1016/j.rser.2012.11.030

Chaudry, S., 2021. Integrating Microalgae Cultivation with Wastewater Treatment: a Peek into Economics. Appl Biochem Biotechnol 193, 3395–3406. https://doi.org/10.1007/s12010-021-03612-x

Chisti, Y., 2007. Biodiesel from microalgae. Biotechnol Adv 25, 294–306. https://doi.org/10.1016/j.biotechadv.2007.02.001

Einhaus, A., Steube, J., Freudenberg, R.A., Barczyk, J., Baier, T., Kruse, O., 2022. Engineering a powerful green cell factory for robust photoautotrophic diterpenoid production. Metab Eng 73, 82–90. https://doi.org/10.1016/j.ymben.2022.06.002

Foladori, P., Petrini, S., Nessenzia, M., Andreottola, G., 2018. Enhanced nitrogen removal and energy saving in a microalgal – bacterial consortium treating real municipal wastewater. Water Science & Technology 78, 174–182. https://doi.org/10.2166/wst.2018.094

Freudenberg, R.A., Wittemeier, L., Einhaus, A., Baier, T., Kruse, O., 2022. Advanced pathway engineering for phototrophic putrescine production. Plant Biotechnol J 1–15. https://doi.org/10.1111/pbi.13879

Gao, F., Yang, Z.Y., Zhao, Q.L., Chen, D.Z., Li, C., Liu, M., Yang, J.S., Liu, J.Z., Ge, Y.M., Chen, J.M., 2021. Mixotrophic cultivation of microalgae coupled with anaerobic hydrolysis for sustainable treatment of municipal wastewater in a hybrid system of anaerobic membrane bioreactor and membrane photobioreactor. Bioresour Technol 337, 125457. https://doi.org/10.1016/j.biortech.2021.125457

Gou, Y., Yang, J., Fang, F., Guo, J., Ma, H., Gou, Y., 2020. Feasibility of using a novel algal-bacterial biofilm reactor for efficient domestic wastewater treatment 3330. https://doi.org/10.1080/09593330.2018.1499812

Gruchattka, E., Kayser, O., 2015. In vivo validation of in silico predicted metabolic engineering strategies in yeast: Disruption of α-ketoglutarate dehydrogenase and expression of ATP-citrate lyase for terpenoid production. PLoS One 10, 1–25. https://doi.org/10.1371/journal.pone.0144981

Gutzeit, G., Lorch, D., Weber, A., Engels, M., Neis, U., 2005. Bioflocculent algal-bacterial biomass improves low-cost wastewater treatment. Water Science and Technology 52, 9–18. https://doi.org/10.2166/wst.2005.0415

Klassen, V., Blifernez-Klassen, O., Bax, J., Kruse, O., 2020. Wastewater-borne microalga Chlamydomonas sp.: A robust chassis for efficient biomass and biomethane production applying low-N cultivation strategy. Bioresour Technol. https://doi.org/10.1016/j.biortech.2020.123825

Lauersen, K.J., 2019. Eukaryotic microalgae as hosts for light-driven heterologous isoprenoid production. Planta 249, 155–180. https://doi.org/10.1007/s00425-018-3048-x

Lauersen, K.J., Baier, T., Wichmann, J., Wördenweber, R., Mussgnug, J.H., Hübner, W., Huser, T., Kruse, O., 2016. Efficient phototrophic production of a high-value sesquiterpenoid from the eukaryotic microalga Chlamydomonas reinhardtii. Metab Eng. https://doi.org/10.1016/j.ymben.2016.07.013

Lauersen, K.J., Wichmann, J., Baier, T., Kampranis, S.C., Pateraki, I., Møller, B.L., Kruse, O., 2018. Phototrophic production of heterologous diterpenoids and a hydroxy-functionalized derivative from Chlamydomonas reinhardtii. Metab Eng 49, 116–127. https://doi.org/10.1016/j.ymben.2018.07.005

Leite, L. de S., Hoffmann, M.T., Daniel, L.A., 2019. Microalgae cultivation for municipal and piggery wastewater treatment in Brazil. Journal of Water Process Engineering. https://doi.org/10.1016/j.jwpe.2019.100821

Li, K., Liu, Q., Fang, F., Luo, R., Lu, Q., Zhou, W., Huo, S., Cheng, P., Liu, J., Addy, M., Chen, P., Chen, D., Ruan, R., 2019. Microalgae-based wastewater treatment for nutrients recovery: A review. Bioresour Technol. https://doi.org/10.1016/j.biortech.2019.121934

Liu, X. ya, Hong, Y., 2021. Microalgae-Based Wastewater Treatment and Recovery with Biomass and Value-Added Products: a Brief Review. Curr Pollut Rep 7, 227–245. https://doi.org/10.1007/s40726-021-00184-6

Mehrshahi, P., Nguyen, G.T.D.T., Gorchs Rovira, A., Sayer, A., Llavero-Pasquina, M., Lim Huei Sin, M., Medcalf, E.J., Mendoza-Ochoa, G.I., Scaife, M.A., Smith, A.G., 2020. Development of Novel Riboswitches for Synthetic Biology in the Green Alga Chlamydomonas. ACS Synth Biol 9, 1406–1417. https://doi.org/10.1021/acssynbio.0c00082

Neupert, J., Gallaher, S.D., Lu, Y., Strenkert, D., Segal, N., Barahimipour, R., Fitz-Gibbon, S.T., Schroda, M., Merchant, S.S., Bock, R., 2020. An epigenetic gene silencing pathway selectively acting on transgenic DNA in the green alga Chlamydomonas. Nat Commun 11. https://doi.org/10.1038/s41467-020-19983-4

Neupert, J., Karcher, D., Bock, R., 2009. Generation of Chlamydomonas strains that efficiently express nuclear transgenes. Plant Journal 57, 1140–1150. https://doi.org/10.1111/j.1365-313X.2008.03746.x

Overmans, S., Ignacz, G., Beke, A.K., Xu, J., Saikaly, P.E., Szekely, G., Lauersen, K.J., 2022. Continuous extraction and concentration of secreted metabolites from engineered microbes using membrane technology. Green Chemistry 24, 5479–5489. https://doi.org/10.1039/d2gc00938b

Overmans, S., Lauersen, K.J., 2022. Biocompatible fluorocarbon liquid underlays for in situ extraction of isoprenoids from microbial cultures. RSC Adv 12, 16632–16639. https://doi.org/10.1039/d2ra01112c

Perozeni, F., Cazzaniga, S., Baier, T., Zanoni, F., Zoccatelli, G., Lauersen, K.J., Wobbe, L., Ballottari, M., 2020. Turning a green alga red: engineering astaxanthin biosynthesis by intragenic pseudogene revival in Chlamydomonas reinhardtii. Plant Biotechnol J 18, 2053–2067. https://doi.org/10.1111/pbi.13364

Qu, W., Loke Show, P., Hasunuma, T., Ho, S.H., 2020. Optimizing real swine wastewater treatment efficiency and carbohydrate productivity of newly microalga Chlamydomonas sp. QWY37 used for cell-displayed bioethanol production. Bioresour Technol 305, 123072. https://doi.org/10.1016/j.biortech.2020.123072

Rajput, B.S., Hai, T.A.P., Gunawan, N.R., Tessman, M., Neelakantan, N., Scofield, G.B., Brizuela, J., Samoylov, A.A., Modi, M., Shepherd, J., Patel, A., Pomeroy, R.S., Pourahmady, N., Mayfield, S.P., Burkart, M.D., 2022. Renewable low viscosity polyester-polyols for biodegradable thermoplastic polyurethanes. J Appl Polym Sci 1–14. https://doi.org/10.1002/app.53062

Ruiz-Martinez, A., Martin Garcia, N., Romero, I., Seco, A., Ferrer, J., 2012. Microalgae cultivation in wastewater: Nutrient removal from anaerobic membrane bioreactor effluent. Bioresour Technol 126, 247–253. https://doi.org/10.1016/j.biortech.2012.09.022

Serna-García, R., Mora-Sánchez, J.F., Sanchis-Perucho, P., Bouzas, A., Seco, A., 2020a. Anaerobic membrane bioreactor (AnMBR) scale-up from laboratory to pilot-scale for microalgae and primary sludge co-digestion: Biological and filtration assessment. Bioresour Technol 316, 123930. https://doi.org/10.1016/j.biortech.2020.123930

Serna-García, R., Zamorano-López, N., Seco, A., Bouzas, A., 2020b. Co-digestion of harvested microalgae and primary sludge in a mesophilic anaerobic membrane bioreactor (AnMBR): Methane potential and microbial diversity. Bioresour Technol 298, 122521. https://doi.org/10.1016/j.biortech.2019.122521

Viruela, A., Murgui, M., Gómez-Gil, T., Durán, F., Robles, Á., Ruano, M.V., Ferrer, J., Seco, A., 2016. Water resource recovery by means of microalgae cultivation in outdoor photobioreactors using the effluent from an anaerobic membrane bioreactor fed with pre-treated sewage. Bioresour Technol 218, 447–454. https://doi.org/10.1016/j.biortech.2016.06.116

Wichmann, J., Baier, T., Wentnagel, E., Lauersen, K.J., Kruse, O., 2018. Tailored carbon partitioning for phototrophic production of (E)-α-bisabolene from the green microalga Chlamydomonas reinhardtii. Metab Eng 45, 211–222. https://doi.org/10.1016/j.ymben.2017.12.010

Wichmann, J., Lauersen, K.J., Biondi, N., Christensen, M., Guerra, T., Hellgardt, K., Kühner, S., Kuronen, M., Lindberg, P., Rösch, C., Yunus, I.S., Jones, P., Lindblad, P., Kruse, O., 2021. Engineering Biocatalytic Solar Fuel Production: The PHOTOFUEL Consortium. Trends Biotechnol 39, 323–327. https://doi.org/10.1016/j.tibtech.2021.01.003

Xiong, Y., Hozic, D., Goncalves, A.L., Simões, M., Hong, P.Y., 2018. Increasing tetracycline concentrations on the performance and communities of mixed microalgae-bacteria photo-bioreactors. Algal Res 29, 249–256. https://doi.org/10.1016/j.algal.2017.11.033

Zhou, Y., He, Y., Zhou, Z., Xiao, X., Wang, M., Chen, B., 2022. A newly isolated microalga Chlamydomonas sp. YC to efficiently remove ammonium nitrogen of rare earth elements wastewater. J Environ Manage 316, 115284. https://doi.org/10.1016/j.jenvman.2022.115284

