## Supplemental files zip for "Biomass generation and heterologous isoprenoid milking from engineered microalgae grown in anaerobic membrane bioreactor effluent": 05 Supp Figs_Waste water manuscript.docx

**SUPPLEMENTARY FIGURES**

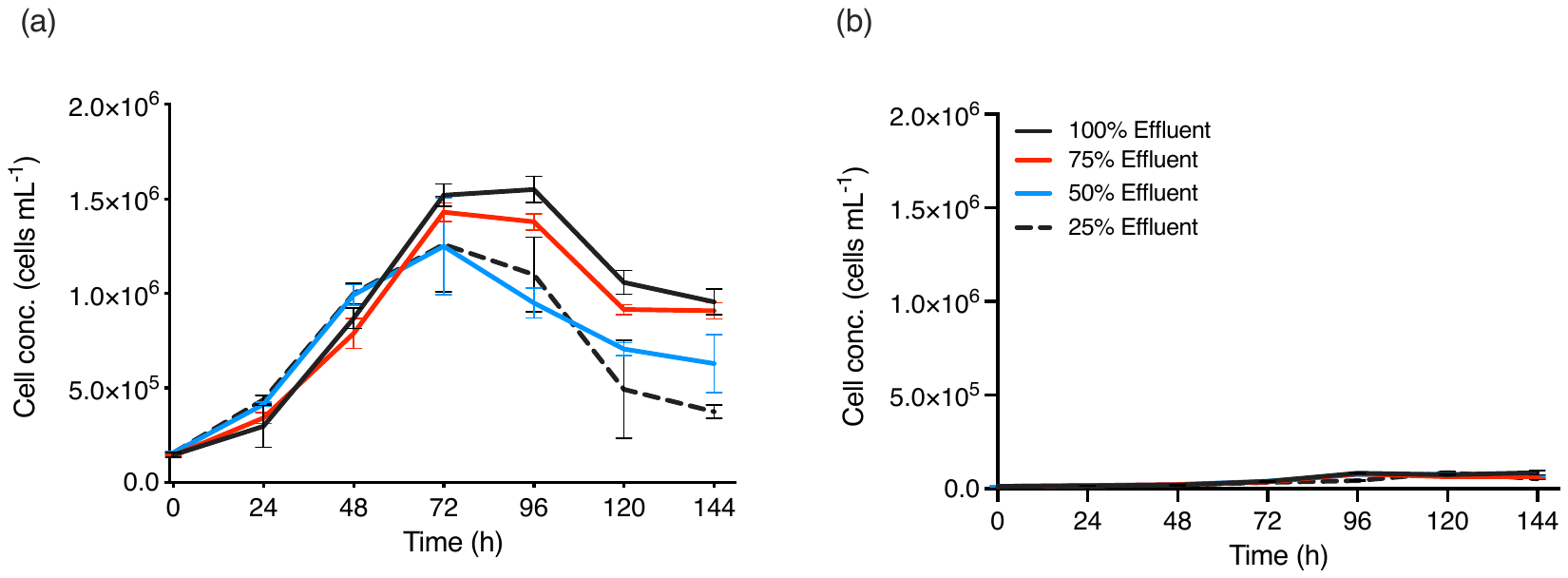

**Suppl. Fig. 1.** Concentrations of (a) *C. reinhardtii* agal cells and (b) bacteria cells during the effluent dilution experiment, where different effluent percentages (25–100%) were used as growth medium.

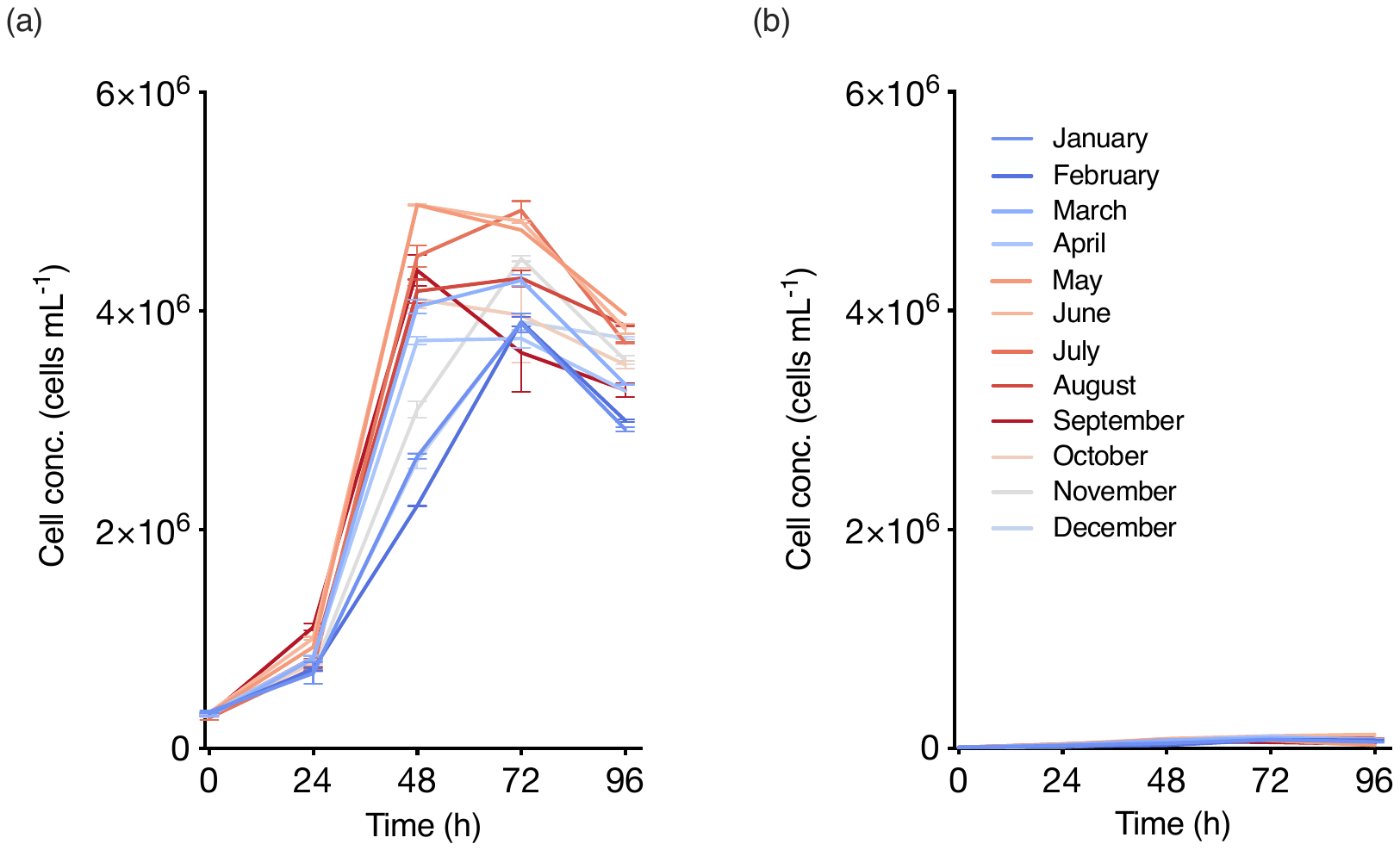

**Suppl. Fig. 2.** Concentrations of (a) *C. reinhardtii* agal cells and (b) bacteria cells during the weather simulation experiment, whereby different incubation conditions based on local temperature and PAR profiles recorded in Thuwal, Saudi Arabia between January and December 2014 were used.

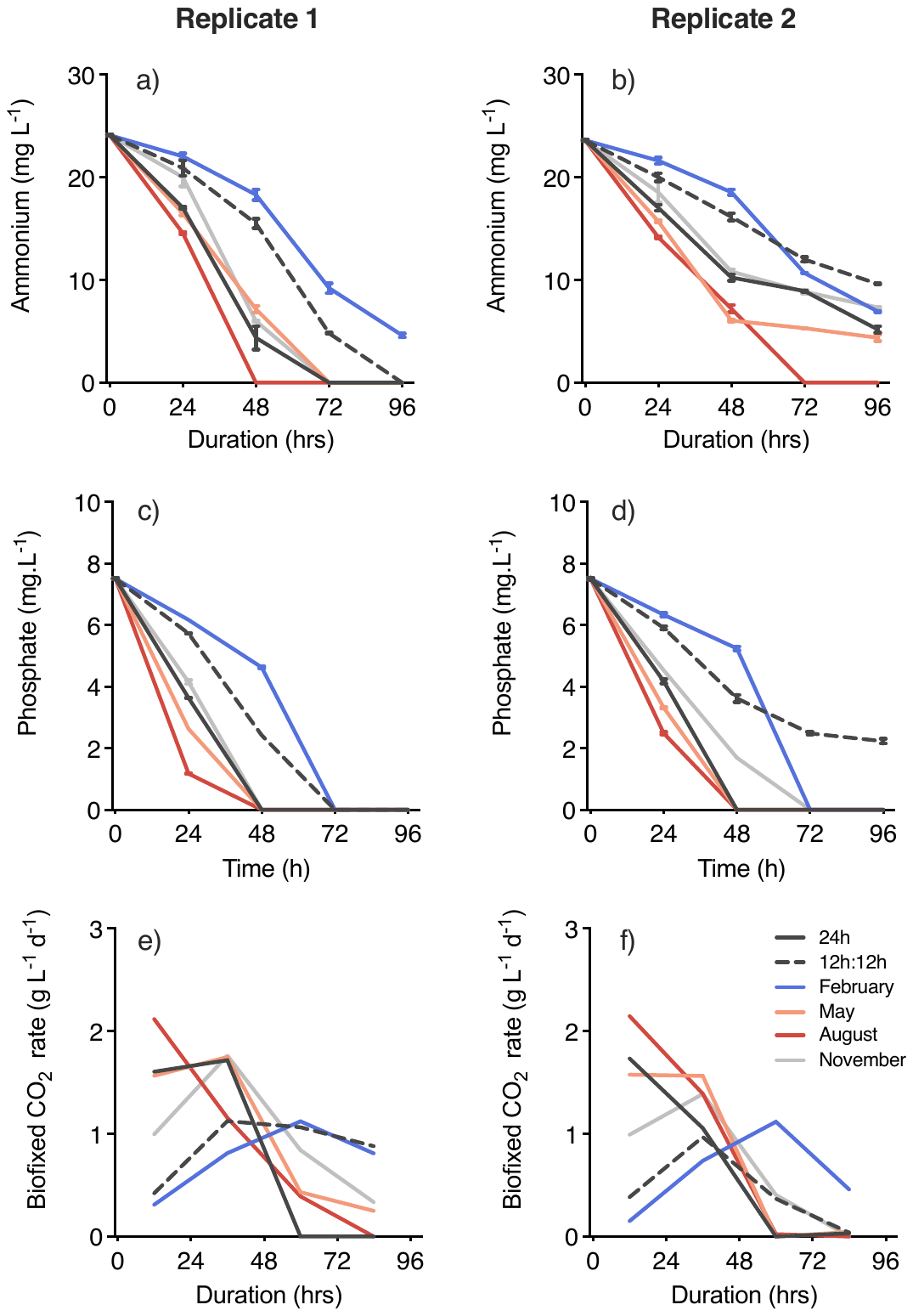

**Suppl. Fig. 3.** (a, b) Ammonium and (c, d) phosphate concentrations and (e, f) bio-fixed CO_2_, calculated as the maximum biomass in a 24 h interval * 1.83 (CO_2_ : biomass ratio) during 96 h of *C. reinhardtii* cultivation in wastewater under six different environmental conditions. Parameters are shown for the second (left column panels) and third (right column panels) experiment. Values displayed in (a)–(d) are means ± SEM of two technical replicates (n=2).

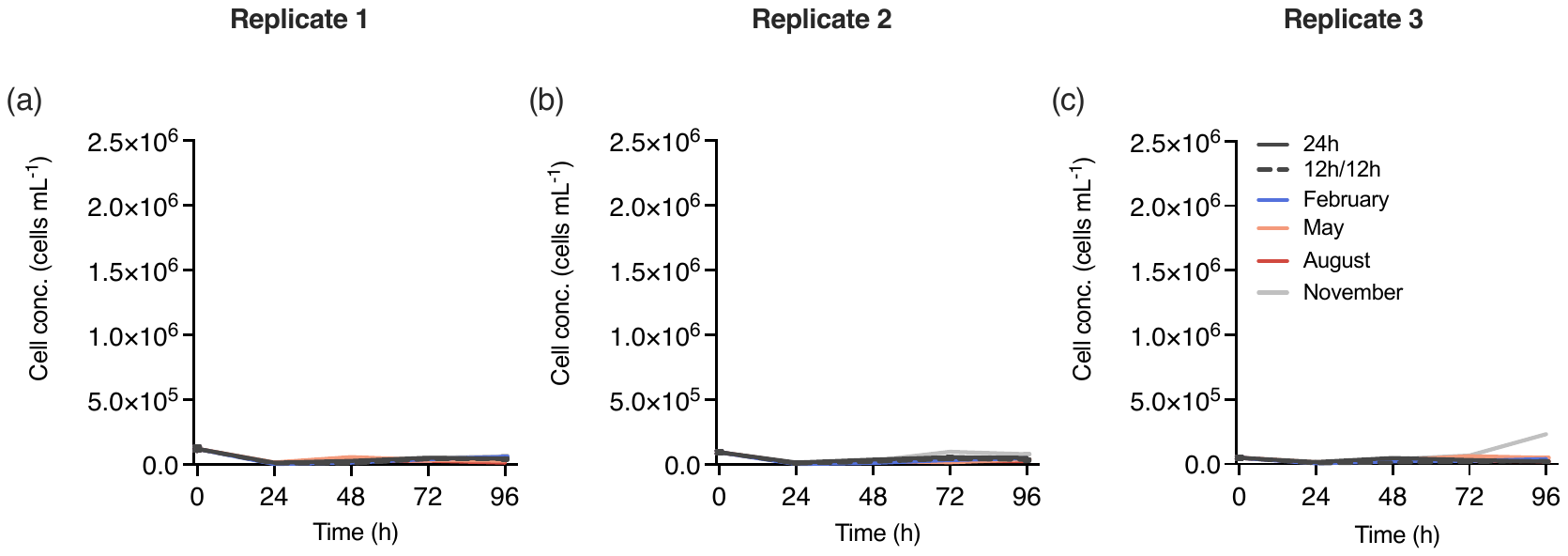

**Suppl. Fig. 4.** Bacterial cell concentrations during 96 h of *C. reinhardtii* cultivation in wastewater under six different environmental conditions. Cell concentrations shown for the first (a), second (b) and third (c) round of experiment.

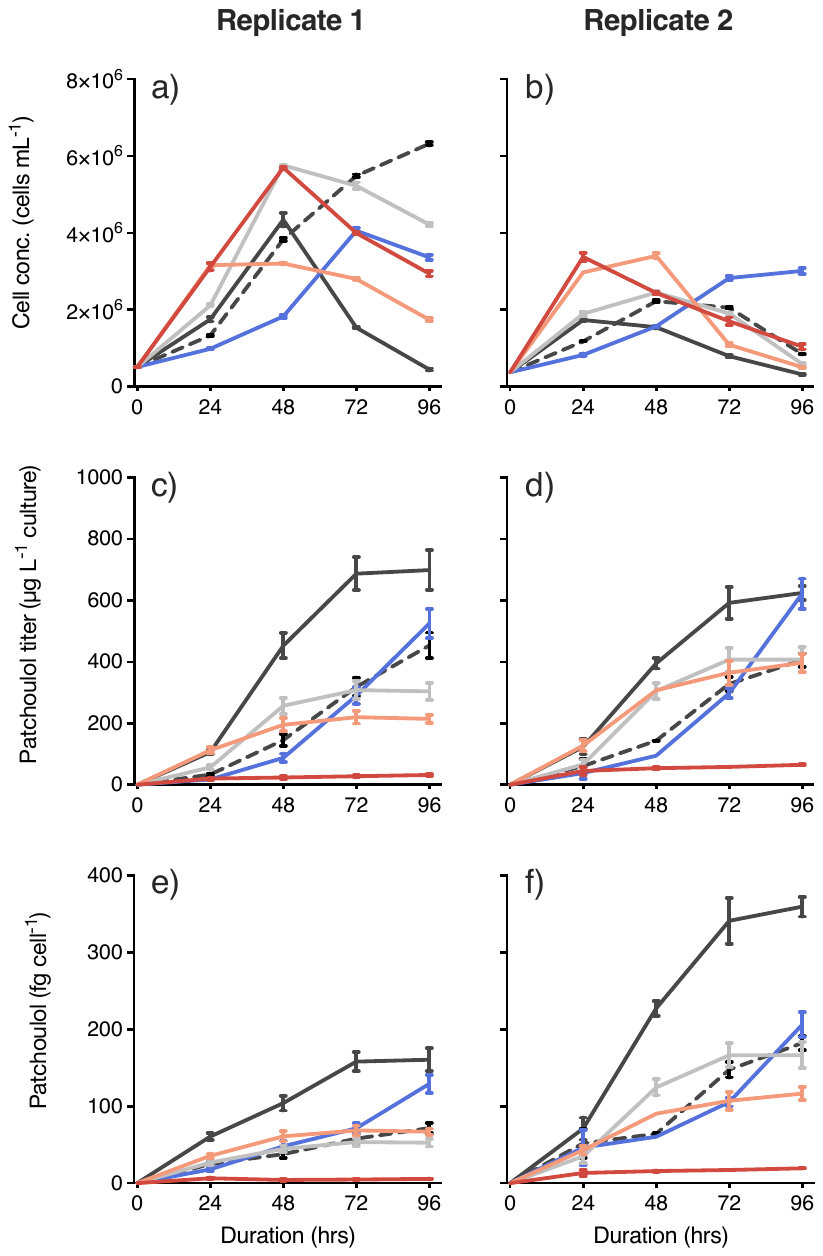

**Suppl. Fig. 5.** (a, b) *C. reinhardtii* algal cell counts during 96 h of cultivation in wastewater in six different environmental conditions. (c, d) Volumetric- and (e, f) cell-dependent patchoulol production during cultivation. Parameters are shown for the first (left column panels) and second (right column panels) experiment. Values displayed are means ± SEM of two technical replicates (n=2).
